# Comparative evaluation on the fermentative potential of single and assorted fruit and vegetable waste for production of bioethanol

**DOI:** 10.1101/2025.07.03.662480

**Authors:** Sudeshna Bera, Sakhi Kundu, Asmita Banerjee, Semantee Bhattacharyya, Sarthak Mondal, Asmita Das, Touhidur Rahaman, Pijus Ghorai, Sukanta De, Jhuma Ganguly, Avishek Banik, Malay Das

## Abstract

Fruit and vegetable waste (FVW) constitute a large part of municipal solid waste (MSW) and create significant environmental challenges by emitting the greenhouse gas methane. In contrast, these are a big source of fermentable sugar to produce bioenergy. Therefore, in this study, the Scanning Electron Microscopy (SEM), Attenuated total reflectance-fourier-transform infrared spectroscopy (ATR-FTIR), thermogravimetric analysis (TGA), X-ray diffraction (XRD), and biochemical quantification of α-cellulose and acid-insoluble lignin contents were carried out to compare fermentative potential between single (potato peel vs. orange peel) and assorted vegetable and fruit wastes. Collectively, the study indicates that assorted vegetable waste holds enormous biofuel potential by having low moisture content (6.78 %), moderately low decomposition temperature (Tmax: 299.36 °C), low crystallinity index (CrI: 19.12%), and high α-cellulose contents (32.35 - 47.73 gm%) yielding high sugar (325.65 mg/g). The study also optimized several key fermentation parameters such as pH, sugar concentration and the time to produce bioethanol. It was observed that maximum production of ethanol (2.14%) was achieved at 28°C, at pH 6.3 with a higher initial sugar concentration of 200mg/g by using *Saccharomyces cerevisiae* strain NCIM 3594. This study supports the likely utility of unsorted vegetable waste to obtain bioethanol at a mass scale.

## 1. Introduction

The stock of non-renewable energy sources is diminishing at a fast pace. In contrast, the need for energy is growing rapidly due to industrialization, particularly in the expanding Asian economies. The International Energy Agency (IEA) predicted that the energy demand of the world would rise by 2.5 times by 2030 [1]. To reduce global warming and the emission of greenhouse gasses, there is an imminent need for a transition from petroleum to renewable energy [2]. Among various renewable energy sources, bioethanol has become quite popular and multiple countries have already adopted Ethanol Blended Petrol (EBP). For instance, the current Indian target of 10% ethanol blending for 2021-22 will be increased to 20% by 2030 (https://mopng.gov.in/en/refining/ethanol-blended-petrol). This is environmentally sustainable and also supports a circular economy. However, the likely success of such a transition from fossil fuel to bioethanol largely depends on the easy availability of biomasses, and their effective conversion to ethanol at a low cost [3, 4].

The global bioethanol industry keeps evolving with the primary goal of affordability, sustainability and quick solution to energy shortage. The first-generation bioethanol was primarily reliant on food or oil-yielding plants such as sugarcane [5], maize [6], and Brassica [7]. However, their massive use in generation of energy has been restricted because of its potential impact on food prices and reduced availability of land for agriculture. Hence, the necessity to move towards non-edible plant biomasses has been prioritized [8]. Therefore, second-generation bioethanol relied on non-crop, non-timber plants such as bamboo [9]; grasses [10, 11] and even crop remains such as- husk [12], and wheat straw [13]. Concomitantly, recycling organic waste obtained from sugarcane bagasse [14]; corn stalk [15]; sorghum stover [16], cassava, ficus fruits [17] and peach waste [18] have also been explored for bioethanol potential. However, limitations such as the requirement of land to grow some of these plants and their unavailability throughout the year are major hindrances behind their wider acceptability for biofuel generation. Utilization of municipal solid waste (MSW) might be an alternative, which could be effective for populated countries like China or India. It had been estimated that Asia’s urban population produces approximately 760 × 103 tons of MSW each day, which is expected to increase to 1.8 × 106 t by 2025 [19]. Another assessment by The Energy and Resources Institute (TERI) projected that India generates over 62 million tons (MT) of waste every year. Only 43 MT of total waste gets collected, out of which 12 MT is treated before disposal, while the remaining 31 MT is simply discarded in waste yards [20]. Regular generation of such a massive amount of MSW poses a serious challenge to solid waste management and disposal in developing countries [21, 22]. In contrast, they are rich sources of fermentable sugar and could be subjected to biorefinery processes to promote a bio-based economy [23, 24, 25]. Particularly, the biogenic MSW is an important, organic source of energy-rich biochemicals comprising carbohydrates, proteins and lipids [26, 27]. Combined use of a set of microorganisms along with exogenous xylanases enabled hydrolysis of both hexose and pentose sugars present in it and resulted in maximum conversion of bio-ethanol [27]. Another study proposed an integrated approach for the maximum use of bagasse obtained from sugarcane by producing bio-H2, and bioethanol along with the utilization of acidogenic effluent for farming [25].

In addition to BMSW, a significant amount of organic waste was generated at domestic or commercial places every day. For instance, approximately 204.83MT of vegetables are annually produced in India every year, out of which ∼12% gets wasted [28]. An older, conservative estimate obtained from the US has predicted that approximately 2030 ±160 trillion BTU of energy was wasted via unused food in 2007. Fruit and vegetable waste (FVW) constitute the major part of organic waste that is regularly being generated in households and markets [29, 30]. For instance, in Europe itself, FVW contributes to approximately 50% of the food waste produced by households regularly [31]. Energy retrieval from FVW can be achieved either via biogas or the generation of bioethanol [32]. Converting FVW to bioethanol is promising for countless reasons, but its commercial viability will depend on the optimization of effective pretreatment, and fermentation processes and their comparative analyses across bio-wastes [33]. It is primarily due to the heterogeneous composition and recalcitrant nature of LCBs, which offers the main impediment in the valorization process. The diversity of LCBs obtained across FVWs might be influenced by seasons, location of collection and economic status of the people. Therefore, the bioenergy potential of individual vegetable or fruit waste should be compared to that of assorted wastes (AW).

The current study aims to compare among individual vegetable, fruit and assorted vegetable waste to transform these bioresources into bioethanol to prevent methane emission and groundwater contamination from decomposing vegetables. Biochemical estimation of α-cellulose and acid-insoluble lignin was combined with analytical parameters (XRD, TGA, FTIR and SEM) to compare the biofuel potential of these biowastes. Further, alkaline pretreatment, saccharification and fermentation reactions were performed to produce ethanol. Our study supports the practical utility of unsorted vegetable waste to obtain bioethanol at the laboratory scale. Further studies are required to evaluate the influence of season, location and other parameters that may influence the yield of bioethanol and the costs accompanying it.

## 2. Material and methods

### 2.1. Selection and collection of vegetable and fruit waste biomasses

One individual vegetable (potato peel, PP), fruit (orange peel, OP) waste along with the assorted vegetable (VW) and fruit wastes (FW) were used for the current analysis (Fig. 1). The detailed composition of assorted vegetable and fruit waste has been provided in Table 1. These were collected from the Koley vegetable market located in Kolkata, India (22.567108, 88.3679842). The vegetable and fruit tissues were dried in a hot air oven overnight at 65°C temperature and were ground by using a mixer grinder to obtain powder to conduct further analyses.

**Fig. 1.**
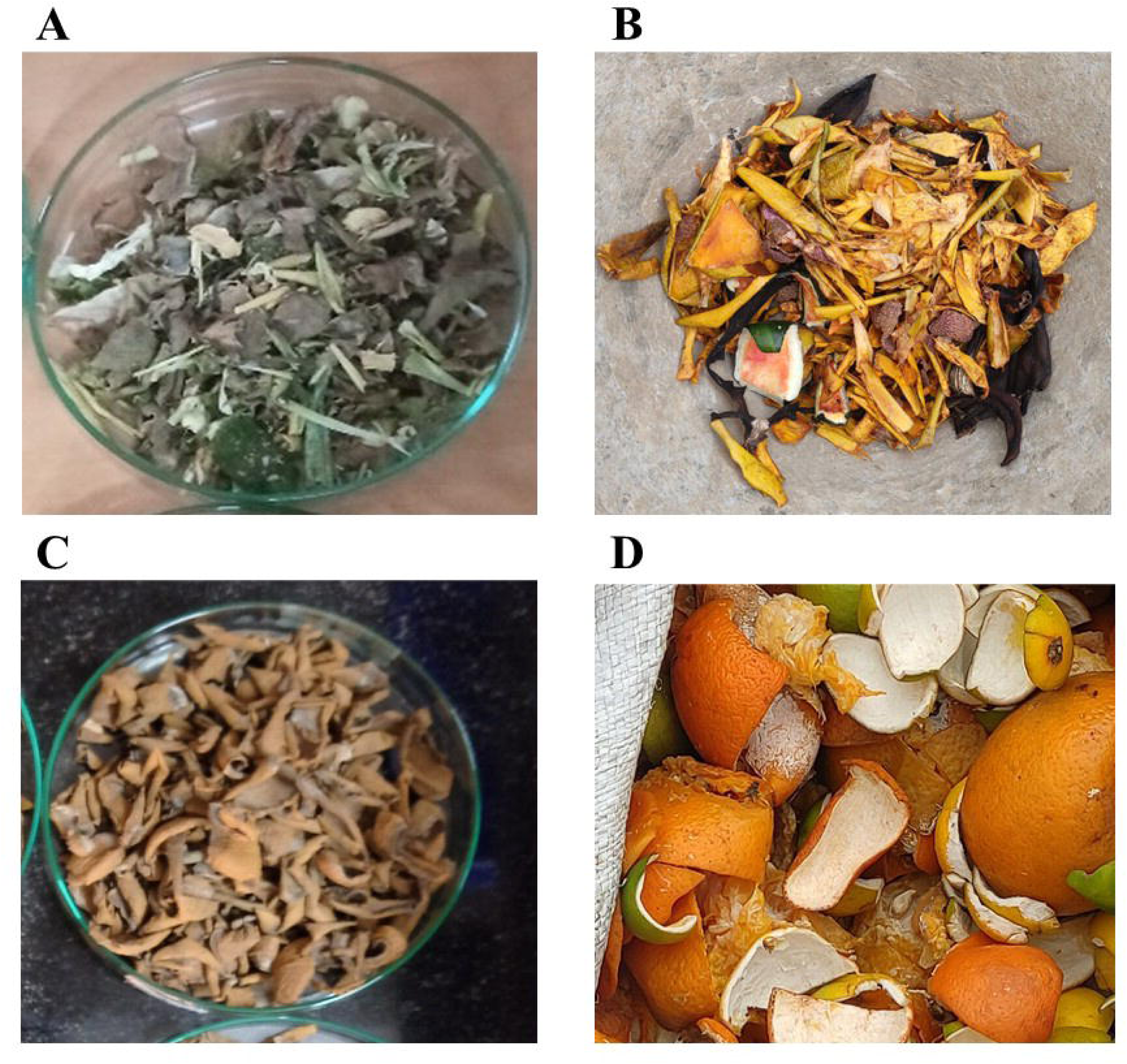
(A-D) Four different FVW tissues used in this study for characterization; mixed vegetable waste (A), mixed fruit waste (B), potato peel (C), Orange waste (D).

**Table 1.** Biochemical estimation of the α-cellulose and acid-insoluble lignin contents in four LCBs.

| LCB Waste | Biological Replicate | Composition | α-cellulose content (gm%) | Acid-insoluble lignin content (gm%) |
| --- | --- | --- | --- | --- |
| Assorted Vegetable waste | Rep 1 | Broccoli, cabbage, cauliflower, pea, coriander, spinach, carrot, radish, unripe papaya, potato peel, kidney-bean | 32.35 | 3.43 |
|  | Rep 2 | Pea, onion, cabbage | 41.43 | 11.45 |
|  | Rep 3 | Wax gourd, Bottle gourd and Pumpkin leaves, pointed gourd, Green Banana, Papaya, Pumpkin, Brinjal, Onion, Lemon, Lady's finger, Bitter gourd (seeds) | 47.73 | 11.98 |
| Assorted Fruit waste | Rep 1 | Orange peel, banana peel, apple peel, jicama, papaya | 28.23 | 6.72 |
|  | Rep 2 | Mango, Banana, Watermelon, Ripe Papaya, Litchi | 15.10 | 4.33 |
|  | Rep 3 | Mango, Banana, Mosambi | 10.72 | 16.6 |
| Orange peel | Rep 1 | - | 18.49 | 2.03 |
|  | Rep 2 | - | 25.33 | 1.70 |
|  | Rep 3 | - | 14.88 | 1.63 |
| Potato Peel | Rep 1 | - | 19.20 | 1.11 |
|  | Rep 2 | - | 15.09 | 8.42 |
|  | Rep 3 | - | 18.19 | 12.08 |

### 2.2. Scanning Electron Microscopy and Attenuated total reflectance-fourier-transform infrared spectroscopy (ATR-FTIR)

To study the effectiveness of pretreatment, 4 unpretreated and pretreated tissues were subjected to scanning electron microscopy (Zeiss EVO10). Approximately, 1 mg of dried powder of PP, OP, AVW and AFW were placed on stubs, mounted with carbon tape and examined at 5kv. Likewise, 1mg of dry, tissue powder was analysed by using ATR-FTIR spectroscopy (Agilent technologies, FTIR-630). Mid-IR spectral range (4000 cm - 400 cm^-1^) with a spectral resolution of 4 cm^-1^ was considered to obtain the spectrogram. Origin Pro software was used to construct an overlapping FTIR spectrogram of all four samples.

### 2.3. Thermogravimetric analysis (TGA)

Thermogravimetry analysis revealed the pyrolytic behavior of samples under investigation. The heating pan of the TG instrument (Discovery 650 SDT thermal analyzer, USA) was loaded with approximately 10 mg of dried, tissue powder. Then samples were heated up to 700 °C under a Nitrogen flow of 50 mL/minute and the heating rate was set at 5 °C/minute. Constant weight loss due to pyrolysis was measured by a mass spectrometer coupled with the TG instrument. Origin Pro software was used to acquire the thermogram (TG) and differential thermogram (DTG). From the thermogram, moisture, hemicellulose and char contents were assessed, while from the DTG curve, maximum thermal degradation (Tmax) was calculated.

### 2.4. X-ray diffraction (XRD)

The Rigaku-Ultima III X-ray diffractometer with Cu Kα radiation (λ = 1.5418 Å) operating at 40 kV, 60 mA was used to analyze the crystallinity of the four studied samples. Scanning range and step size were kept at 2θ =7-40° and 0.02°, respectively. After preparing the diffractogram in Origin Pro software, amorphous and crystalline peaks of cellulose were identified. Segal’s peak height method was utilized for the measurement of the Crystallinity index (CrI, [34].

### 2.5. Measurement of α-cellulose and acid-insoluble lignin

The α-cellulose present in the plant cell wall, can be digested to produce the hexose sugar (glucose) that can be fermented by many yeast strains. Hence biochemical contents of α-cellulose were assessed by the procedure, which was effective to other plant biomasses as well [9, 11]. Approximately, 500 mg of tissue powders were taken, mixed with 1.5 mL nitric acid: acetic acid (1:8 v/v) solution and incubated for 30 minutes in a boiling hot water bath. Removal of hemicellulose and lignin was achieved by repeated washing of the pellet and centrifugation at 6000 rpm until the solution became colorless. Then the pellet was dissolved in 5 mL of 67% sulphuric acid, and kept overnight at 4^0^C followed by mixing with 3 mL of 0.2% chilled anthrone to obtain bluish green colour (SRL, India). The absorbance was measured at 620 nm wavelength in a UV-VIS spectrophotometer (Shimadzu 2600, Japan). The quantity of α-cellulose was calculated as a gram percentage of dry weight by plotting it against a standard curve for cellulose (SRL, India).

Lignin hinders the activity of cellulolytic enzymes during saccharification reaction and acid-insoluble lignin (AIL) is the most abundant lignin present in the cell wall. Hence estimation of AIL was done by following a protocol of the American Society for Testing and Materials [ASTM E1721-01(2015)] as reported earlier [11].

### 2.6. Alkaline pre-treatment, saccharification and measurement of glucose by the Glucose Oxidase- Peroxidase (GOD-POD) method

Removal of lignin is necessary for the saccharification reaction to release fermentable sugar from cellulose. Therefore, chemical pretreatment was undertaken using a 2% NaOH solution due to its wide applicability on plant lignocelluloses [24]. This reaction was carried out at 100 °C for 60 minutes in a dry furnace. To remove alkali, the treated tissues were repeatedly washed with distilled water until the pH of the effluent reached at 7. The pretreated tissues were dried at 60 °C overnight before saccharification. For enzymatic saccharification, 1% (w/v) pretreated tissues were mixed with cellulase enzyme obtained from *Trichoderma reesei* (Sigma, 60unit/g- solid cellulase loading) in 0.05 M citrate buffer (pH- 4.8) and incubated at 50°C with rotational speed of 150 rpm. Three replicates were used for each tissue and aliquots were taken at 8, 24, 48 and 72h post-incubation. In order to measure glucose yield, the GOD-POD method (Autospan®, ARKRAY, India) was used [11]. Detection of absorbance was done at 505 nm in a UV-VIS spectrophotometer (Shimadzu 2600, Japan). Production of glucose was also confirmed by observing the diagnostic mass fragmentation pattern in the High-Resolution Mass Spectrometry (HRMS) analysis (Supplementary Fig. 1).

### 2.7. Effects of pH, incubation time and glucose loading on ethanol yield

Fermentation experiments were performed using yeast (*Saccharomyces cerevisiae* strain NCIM 3594) procured from the National Collection of Industrial Microorganisms (https://www.ncl-india.org/files/ncim/Default.aspx). The strain was cultured in liquid MGYP (malt extract: 3g/L, glucose: 10g/L, yeast extract: 3g/L, and peptone: 5g/L) media overnight, followed by growth in MYGP-agar to obtain a single isolated colony. One mL of yeast inoculum (OD- 0.7) was mixed with 19 mL of enzymatic hydrolysate obtained at 48 hours post hydrolysis. The fermentation reaction was carried out at 28^0^C, 150 rpm for 48 hours in an incubator shaker [35]. Finally, ethanol yield was calculated by using the Potassium dichromate method [36]. Chromium ions present in potassium dichromate reacts to ethanol to produce blue-green colour and the absorbance was measured at 595 nm. The yield of ethanol was estimated using a standard curve prepared using a dilution series of absolute ethanol (0.2-6%) and was expressed in % (v/v). The impact of pH and concentration of initial sugar on ethanol yield was evaluated. Four pH (6, 6.3, 6.6, 6.9) and four initial sugar concentration (50, 100, 150, 200 mg/g) was used. Also, in order to study glucose utilization by yeast, comparison between consumption of glucose vs. ethanol yield from VW and PP biomasses at six specific time points (12, 24, 36, 48, 60, 72h) was conducted.

## 3. Results

### 3.1. Efficacy of pretreatment by SEM analyses

Scanning electron microscopy was carried out to compare and evaluate changes in structural integrity between unpretreated and pretreated tissues. Indeed, unporous and compact tissue structures were observed for the unpretreated biomasses. In contrast, the disintegration of the cell wall matrix and an increase in surface porosity were observed in alkaline pretreated tissues (Fig. 2A-H). This is possibly due to the breakdown of hydrogen bonds in the cellulose chain, which results in the development of porous structures.

**Fig. 2.**
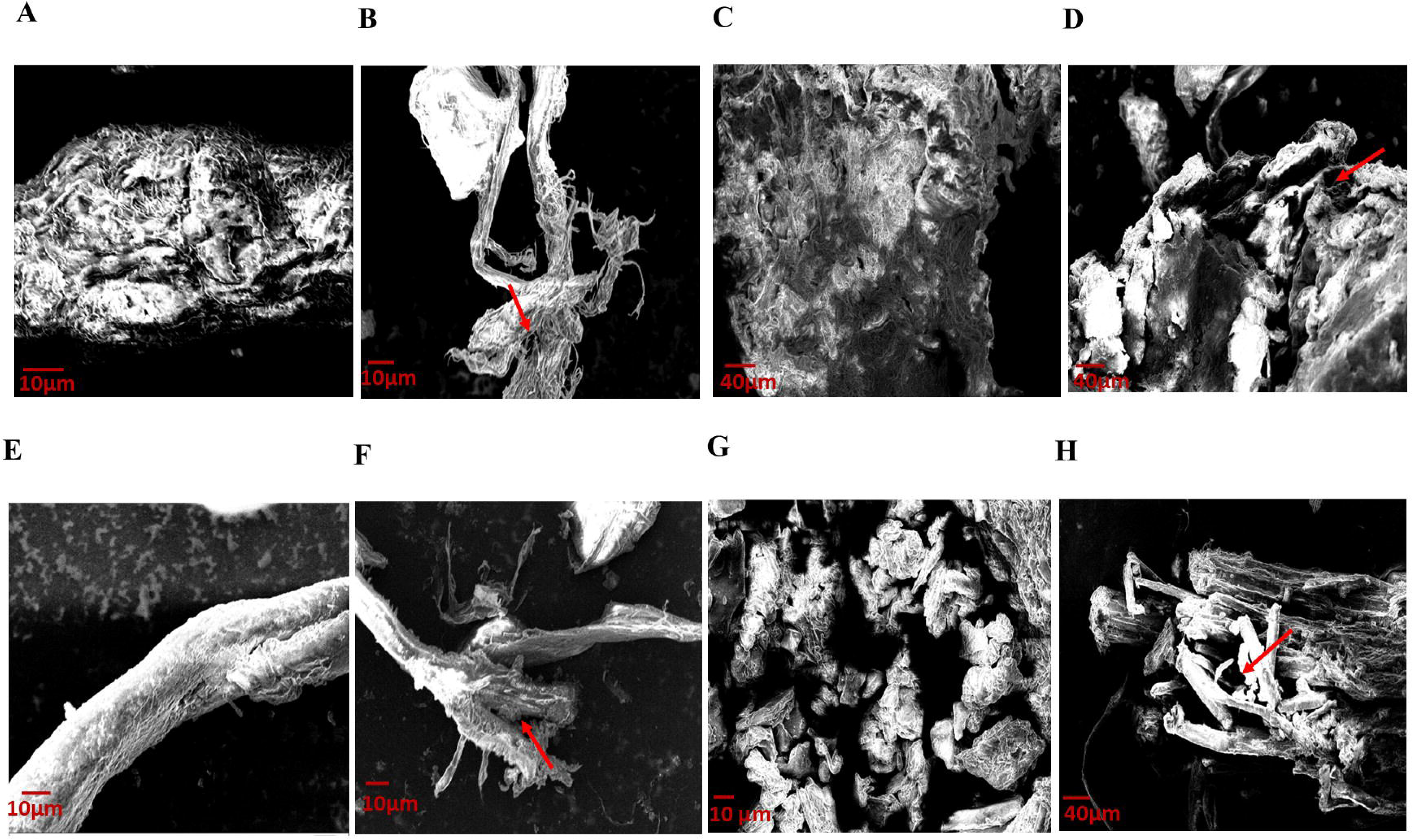
(A-H) Scanning electron microscopy images of untreated vegetable waste (A), potato peel (B), mixed fruit waste (C), orange peel (D) and pretreated vegetable waste (E), potato peel (F), mixed fruit waste (G), orange peel (H). Red arrows indicating disintegrated cell wall matrix due to the alkaline pre-treatment reaction.

### 3.2. Study of the chemical composition of PP, OP, VW and FW by ATR- FTIR

Qualitative assessment of the cell wall constituents of vegetable and fruit wastes were done by the ATR-FTIR spectroscopy analyses. Signature IR peaks, which are representative of major compounds were identified by comparing information from published articles [11, 37, 38, 39, 40, 41]. Spectrogram of untreated and pretreated tissues were compared to assess the effectiveness of the pretreatment reaction. Indeed, variation in peak intensity, addition of new and elimination of older peaks were observed in pretreated samples. The abundance of principal cell wall constituents like cellulose, hemicellulose and lignin was evident from the spectrograms.

Cellulose, hemicellulose, and lignin all exhibit few common peaks according to FTIR analysis, a broad peak at ∼3291 cm⁻¹ that represents O–H stretching, a peak at ∼2913 cm⁻¹ that is due to C–H stretching of aliphatic CH2 groups, and a peak at ∼1004 cm⁻¹ that represents C–O stretching (Table 2, Fig. 3). However, lignin exhibits a unique peak at ∼1601 cm⁻¹, which is caused by aromatic C=C stretching, while hemicellulose is distinguished by a peak at ∼1730 cm⁻¹, which is caused by C=O stretching (Table 2, Fig. 3). The stronger ∼1730 cm⁻¹ peaks in the untreated samples (VW, PP, FW, and OP) show that the FW and OP samples have a higher hemicellulose content than the VW and PP samples. The IR spectra show a significant drop in the ∼1730 cm⁻¹ peak in all samples after pretreatment (majorly visible in P-FW and P-OP as this peak lies prominently), indicating a decrease in the amount of hemicellulose with respect to the untreated samples (Fig. 3). In the case of lignin, in P-FW and P-OP, the intensity of the ∼1601 cm⁻¹ peak drops, suggesting a decrease in lignin content. P-VW and P-PP, on the other hand, appear to have increased in this peak. This could instead be a relative intensification brought on by the removal of hemicellulose, which makes the lignin peak more noticeable, rather than a real increase in lignin concentration. In conclusion, the FTIR data confirm a clear reduction of hemicellulose across all pretreated samples with respect to the untreated samples. The decrease in lignin content is evident for P-FW and P- OP, but less conclusive for P-VW and P-PP due to overlapping spectral effects.

**Fig. 3.**
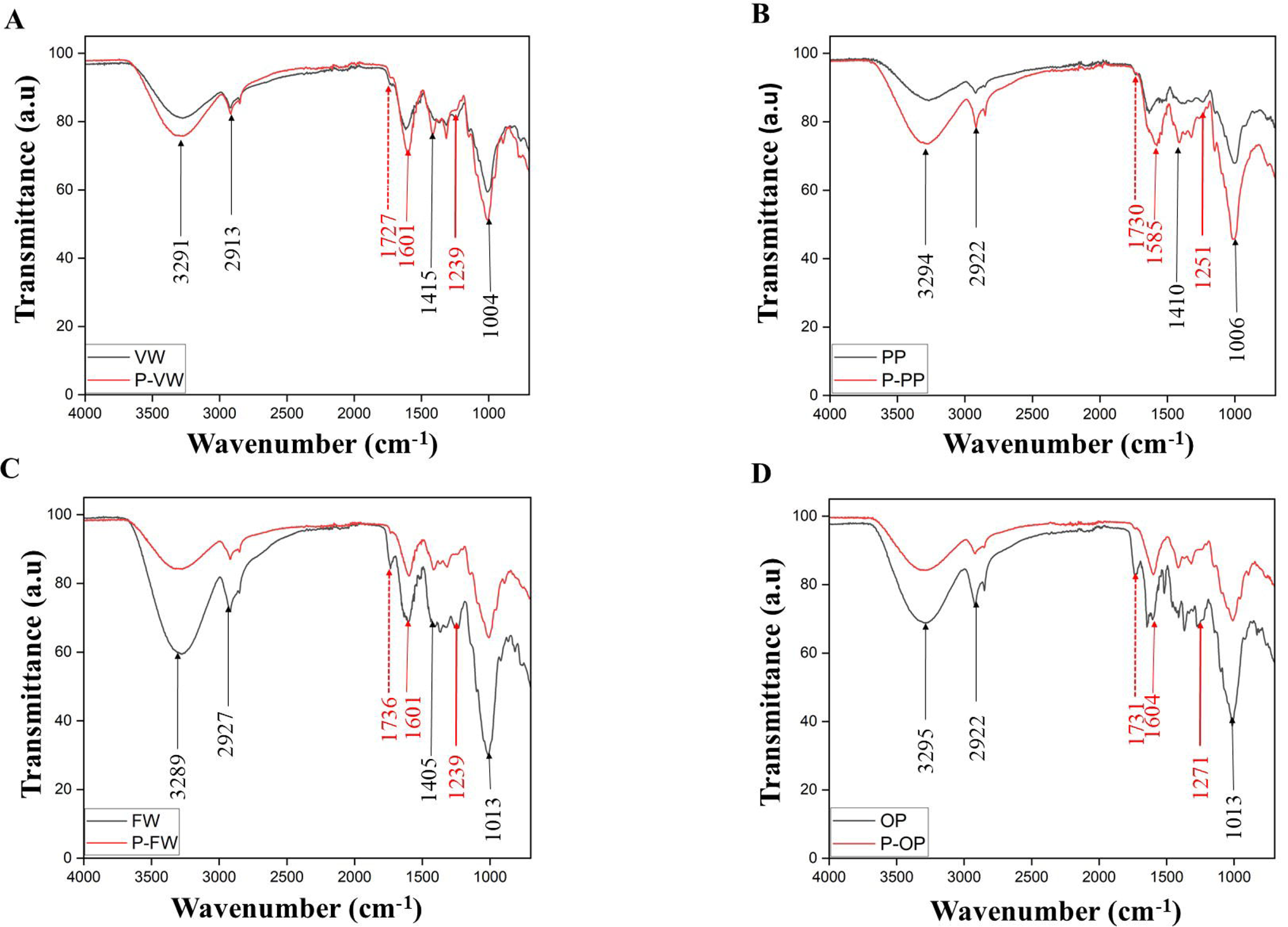
(A-D) Overlaid ATR-FTIR spectrogram of four un-treated and pre-treated FVW tissues- VW (A), PP (B), FW (C) and OP (D). Abbreviations used: VW- vegetative waste, PP- potato peel, FW- fruit waste, OP- orange peel; PVW- pretreated vegetable waste, PPP- pretreated potato peel, PFW- pretreated fruit waste, POP- pretreated orange peel. The black arrow indicates cellulose, the red arrow indicates lignin and the red dotted arrow indicates hemicellulose specific peaks.

**Table 2.** FTIR Comparison of Cellulose, Hemicellulose, and Lignin.

| **Wavenumber (cm<sup>-1</sup>)** | **Functional Group** | **Cellulose** | **Hemicellulose** | **Lignin** |
| --- | --- | --- | --- | --- |
| ~3291 | O–H stretching (hydrogen bonding) | Broad, strong | Broad, slightly less intense | Broad, due to phenolic & alcoholic OH |
| ~2913 | C–H stretching (aliphatic CH<sub>2</sub>) | Present | Present | Present |
| ~1727 | C=O stretching (ester/acetyl) | Absent | Present (acetyl, uronic esters) | Weak or absent |
| ~1601 | Aromatic C=C stretching | Absent | Absent | Strong, characteristic of aromatics |
| ~1415 | CH<sub>2</sub> bending / crystallinity marker | Strong | Weak | Weak or absent |
| ~1319 | C–H bending | Present | Present | Weak |
| ~1239 | C–O stretching (ether, phenol) | Weak or absent | Present (minor) | Strong (aryl ethers, phenols) |
| ~1004 | C–O stretching | Strong | Present | Present (guaiacyl/syringyl units) |

### 3.3. Comparison of thermal stability and estimation of moisture, hemicellulose and residual char content by Thermogravimetry

Pyrolysis of four studied FVWS were performed in an anoxygenic atmosphere to assess their thermal stability, moisture, hemicellulose and char contents. The TGA curve showed a four-step degradation of these tissues. In the first step, weight loss was caused due to evaporation of moisture between 100-150 °C temperature (Fig. 4A). Moisture content in analyzed samples varied from 6.5% in PP to 11.77% in OP. In the second phase, hemicellulose degradation took place between 250-300 °C of temperature. Weight loss due to hemicellulose decomposition ranged from 25.32% in VW to 32.43% in PP (Table 3). In the third step of pyrolysis (300-350 °C), the primarily breakdown of cellulose occurred, which caused a sharp drop in weight. After that, slow degradation of lignin took place until 700 °C and eventually the residual char was left. Char content was ranged from 20.8% in OP to 30.2% in VW. From the DTG curve, the maximum decomposition temperature (Tmax) was estimated for the studied samples, which provided information on their thermal stability (Fig. 4B). Biomass having less thermal stability is usually favored for bio-energy production. In this study, PP had the lowest Tmax (281.66 °C), which was followed by VW (Tmax: 299.36 °C), while OP had the highest Tmax (317.66 °C). Hence, PP and VW might be suitable for bio-energy purposes since they contain low Tmax and moisture content.

**Fig. 4.**
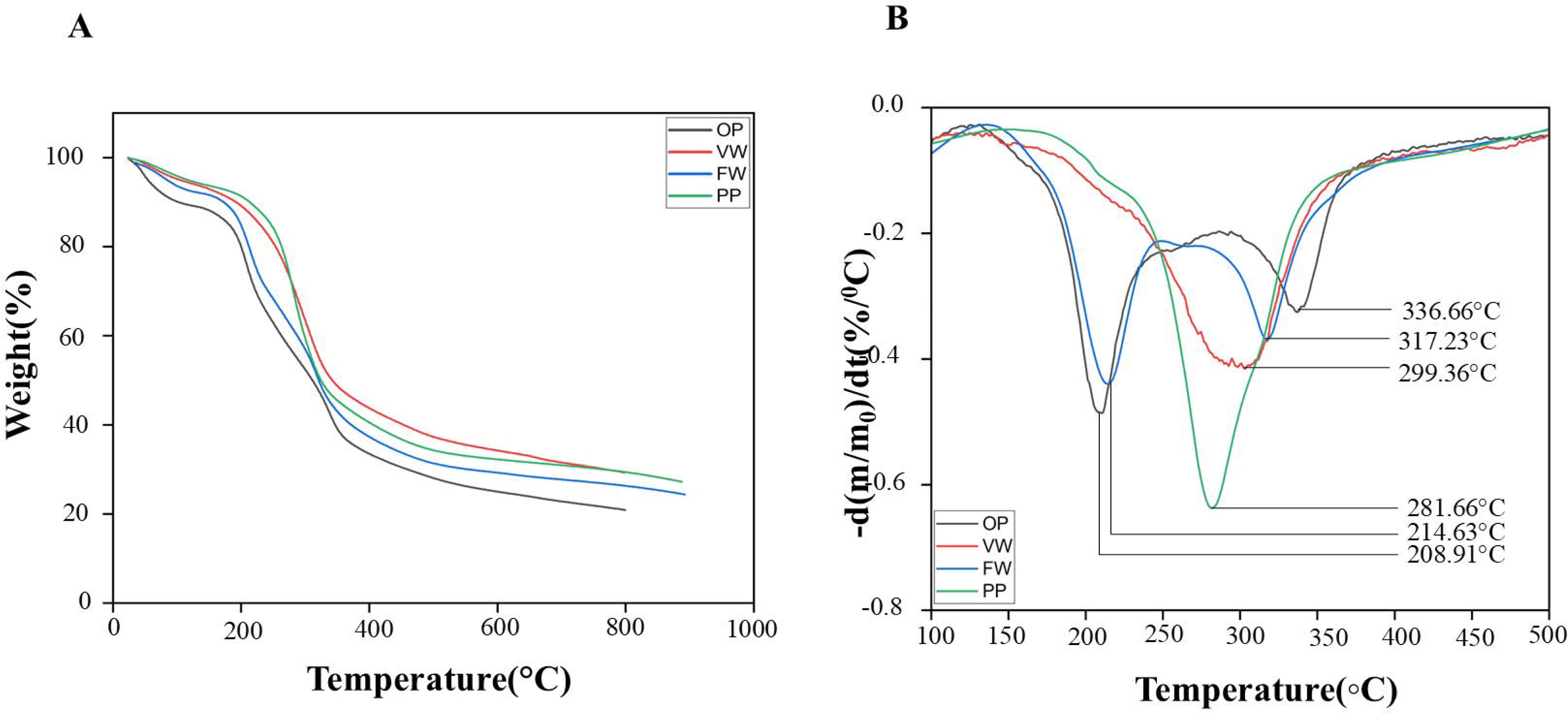
(A-B) Thermogram of four FVW tissues showing their pyrolytic behavior (A) and differential thermogram of the TG curve (B) showing maximum decomposition temperature (Tmax) of the four waste tissues.

**Table 3.** Summary of important tissue parameters deduced from TGA profile of four FVWs.

| **Sample** | **Moisture**<br>**content**<br>**(%)** | **Hemicellulose (%)** | **Residual char (%)** | **Maximum**<br>**decomposition**<br>**temperature**<br>**(T<sub>max</sub>, °C)** |
| --- | --- | --- | --- | --- |
| Assorted<br>Vegetable waste | 6.78 | 25.32 | 30.2 | 299.36 |
| Potato peel | 6.5 | 32.43 | 27.2 | 281.66 |
| Fruit waste | 8.5 | 27.78 | 24.43 | 317.23 |
| Orange peel | 11.77 | 29.15 | 20.8 | 336.66 |

### 3.4. Comparison of cellulose crystallinity by X-ray diffraction

The degree of crystallinity of LCBs influence the success of enzymatic hydrolysis. During saccharification, cellulases are less effective on crystalline cellulose than amorphous ones. Hence, cellulose fibers having a low crystallinity index (CrI) are desirable for bioethanol production. For the determination of CrI, two major peaks were considered. One was I002 (peak for crystalline cellulose) and the other one was IAM (peak for amorphous cellulose). The I002 peak for the studied samples were at 21.5 °2θ - 21.7 °2θ angle and IAM peak was at 17.3° 2θ - 17.4° 2θ angle (Table 4, Fig. 5). The VW had the lowest CrI (19.12%), which was followed by PP (19.76%). These two tissues are expected to demonstrate enhanced efficiency toward an enzymatic breakdown of cellulose. In contrast, FW had the highest CrI (24.59%), followed by OP (22.18%) and hence might be less reactive to enzymatic saccharification.

**Fig. 5.**
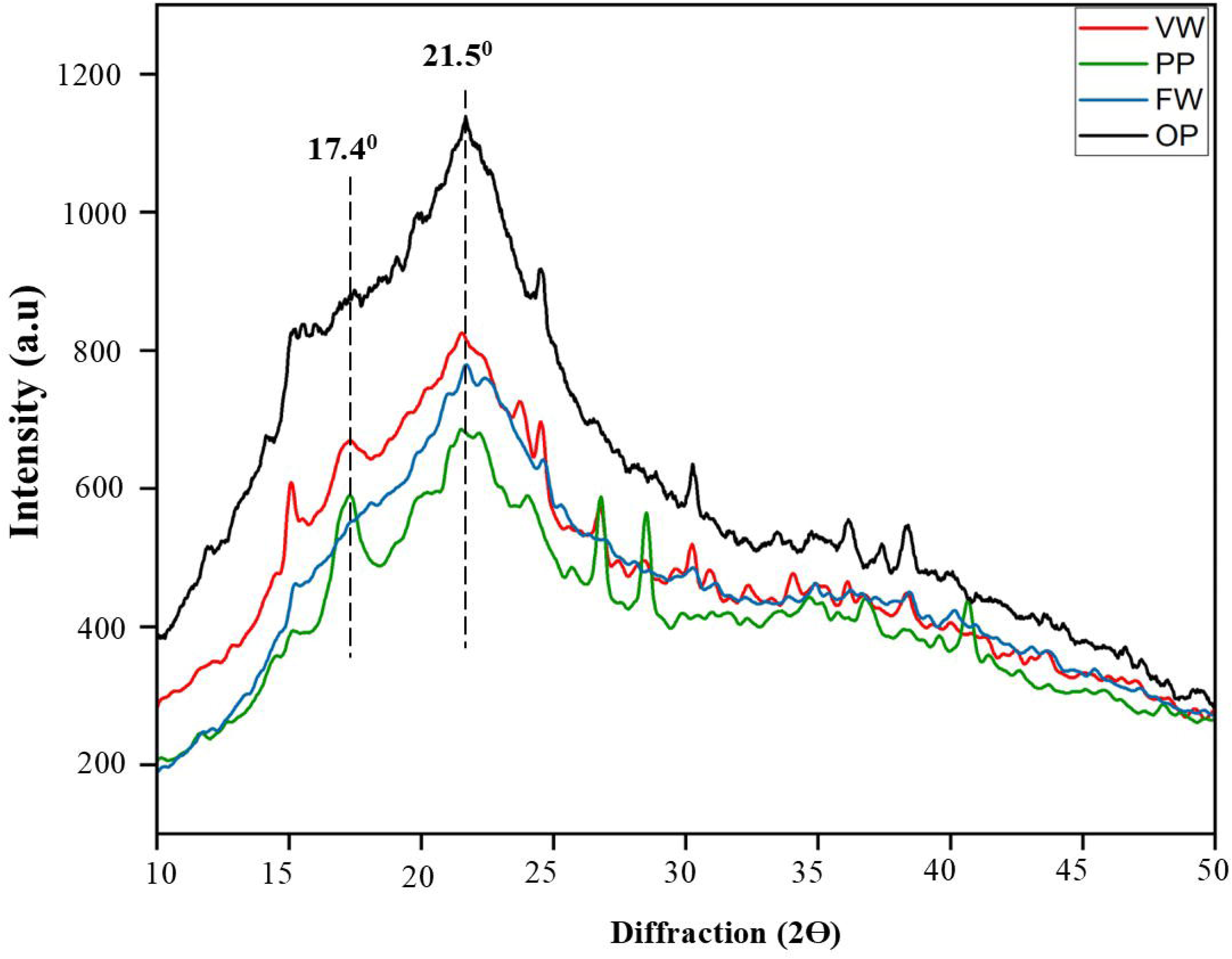
X-ray Diffraction patterns of four FVW tissues displaying peaks for the amorphous and crystalline cellulose. Abbreviations used: VW- vegetable waste, PP- potato peel, FW- fruit waste, OP- orange peel.

**Table 4.**
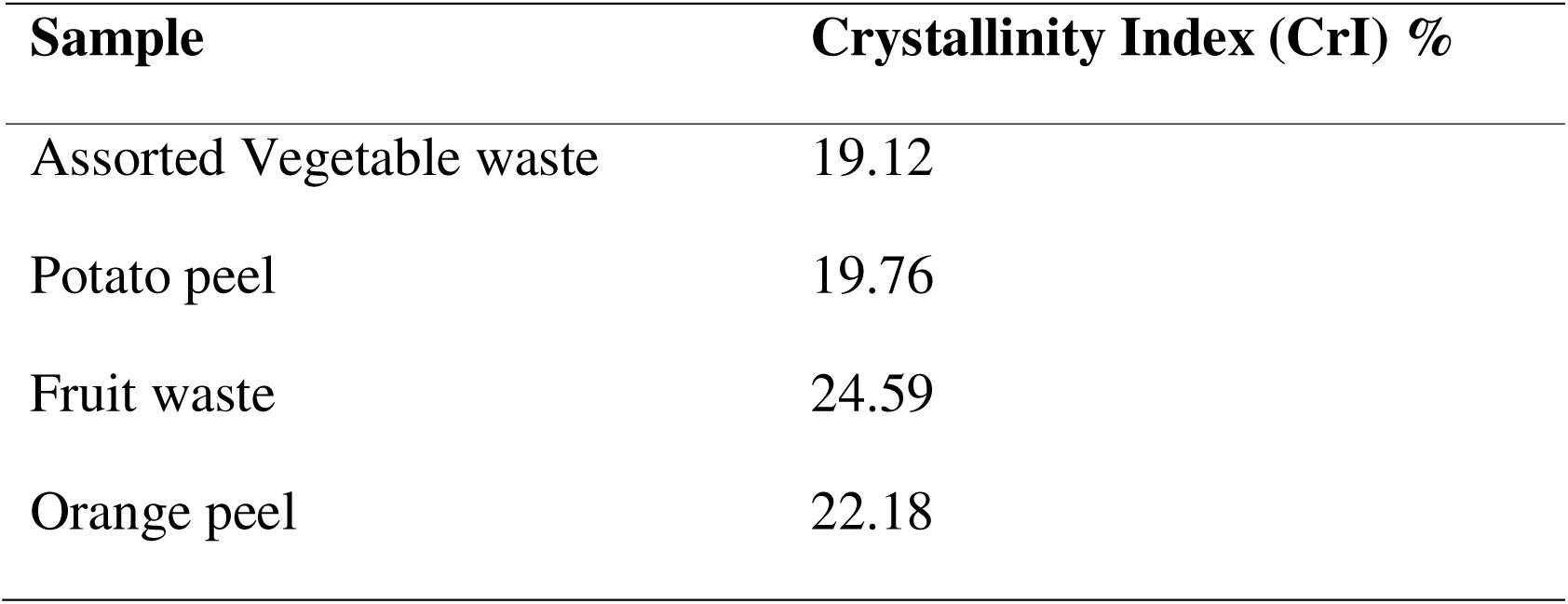
Comparison of cellulose crystallinity index of four FVW tissues.

| **Sample** | **Crystallinity Index (CrI) %** |
| --- | --- |
| Assorted Vegetable waste | 19.12 |
| Potato peel | 19.76 |
| Fruit waste | 24.59 |
| Orange peel | 22.18 |

### 3.5. Biochemical comparison of α-cellulose and acid-insoluble lignin contents

A total of 72 samples (4 samples X 3 biological replicates X 3 technical replicates) obtained from 4 FVW tissues were subjected to biochemical measurements of α-cellulose and acid-insoluble lignin (AIL) contents. They demonstrated a large variation concerning both α-cellulose and AIL levels. Among four tissues, VW represented the highest amount of α-cellulose, which varied from 32.35 - 47.73 gm% (Table 1). Biochemical α-cellulose contents detected in the other three tissues were much lower. It was 10.72 - 28.23 gm% in FW, 14.88 - 25.33 gm% in OP and 15.09 - 19.20 gm% in PP (Table 1). In terms of variation across bio-replicates, PP was the most consistent.

Similarly, the AIL levels demonstrated considerably less variation not only among tissues but also across bio-replicates of one particular tissue. For instance, two replicates of VW contained a moderate amount of lignin (11.45, 11.98 gm%, Table 1), while one replicate contained a much lesser amount (3.43 gm%, Table 1). Similarly, AIL contents ranged from 4.33 - 16.6 gm% in FW, and 1.11 - 12.08 gm% in PP. In contrast, lesser variation was observed for OP (1.63 - 2.03 gm%).

### 3.6. Comparison of glucose yield due to enzymatic hydrolysis

The quantity of fermentable sugar released after saccharification is an important index to assess the bioenergy prospect of a tissue. Therefore, glucose levels were quantified in PP, OP, VW and FW tissues at 6, 24, 48 and 72 h post-saccharification time points (Fig. 6). Kinetic change in sugar yield was very prominent in PP, OP, and VW, but less apparent in FW. For all the tissues, the maximum upsurge in yield was observed between 24 - 48h, after which a more or less steady state was achieved. At a 48h time-point, the yield of glucose was much higher in VW (325.65 mg/g) and PP (317.17 mg/g) than OP (302.98 mg/g) and FW (247.57 mg/g). The high-resolution mass spectrometry analysis at 48h post saccharification time-point detected 3 peaks (205, 343 and 161), that are confirmatory to the generation of the hexose sugar (Supplementary Fig S1).

**Fig. 6.**
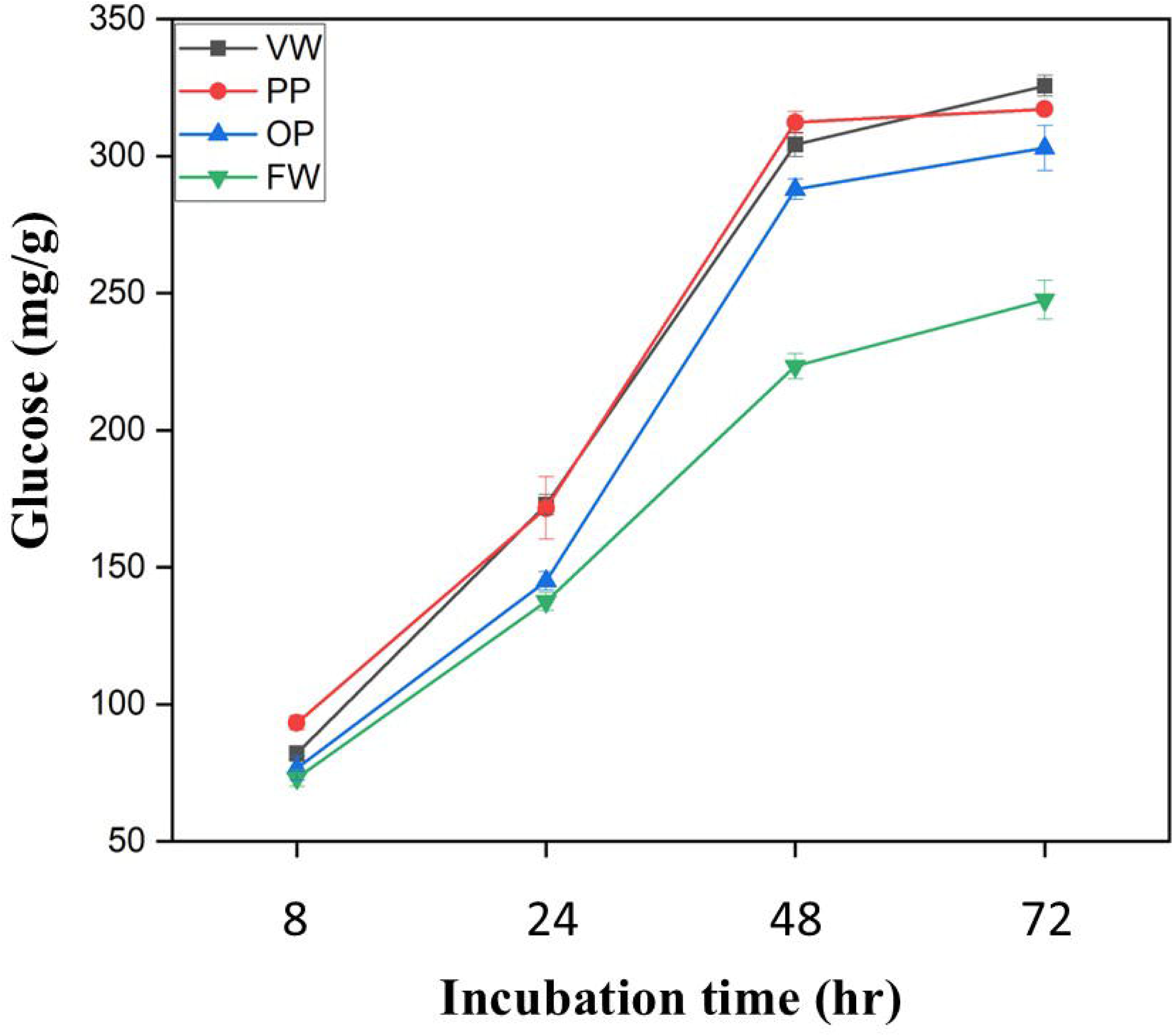
Yield of glucose assessed from four pretreated FVW tissues after 72 hours of enzymatic hydrolysis. Abbreviations used: FW- fruit waste, OP- orange peel, PP- potato peel, VW- vegetable waste.

### 3.7. Comparison of ethanol production after fermentation

Ethanol quantification was conducted on VW and PP since they demonstrated higher sugar yield than the other two FVWs at 48 hours of post-fermentation. Fermentation reaction was conducted at four different pH regimes (6, 6.3, 6.6, 6.9) to identify the optimum pH. Maximum ethanol yield was observed at pH 6.3 at 60 hours of post-fermentation for VW (2.03%), which was followed by PP (1.74%). To identify the impact of sugar loading on ethanol yield, the starting sugar concentration (50, 100, 150, 200 mg/g) was varied, while the pH remained constant at 6.3. The yield of obtained ethanol was increased with the rise of the starting sugar concentration (Fig. 7, Table S1). For instance, the ethanol yield from 50 mg/g loaded sugar was ∼0.55% at 60h post-fermentation for VW, which went up to 2.06% on using 200mg/g of initial sugar concentration. When glucose utilization vs. ethanol yield was compared, amount of remaining glucose was reduced with the progression of time, while ethanol yield was elevated until 60h (Fig. 8).

**Fig. 7.**
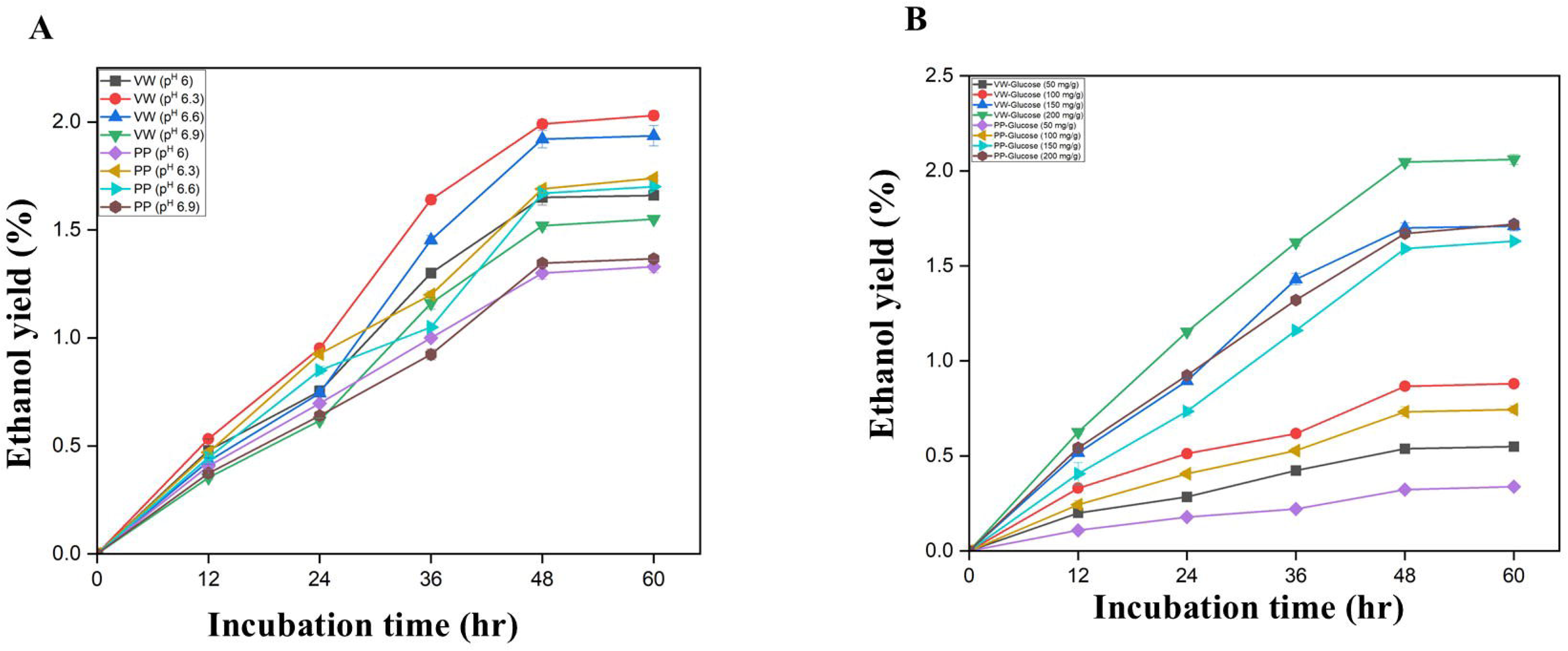
(A, **B)** Assessing impact of pH (6, 6.3, 6.6, 6.9, A) and glucose loading (50, 100, 150, 200 mg/g, B) on ethanol yield at multiple time points (12, 24, 36, 48, 60h) from VW and PP using *Saccharomyces cerevisiae* strain NCIM 3594.

**Fig. 8.**
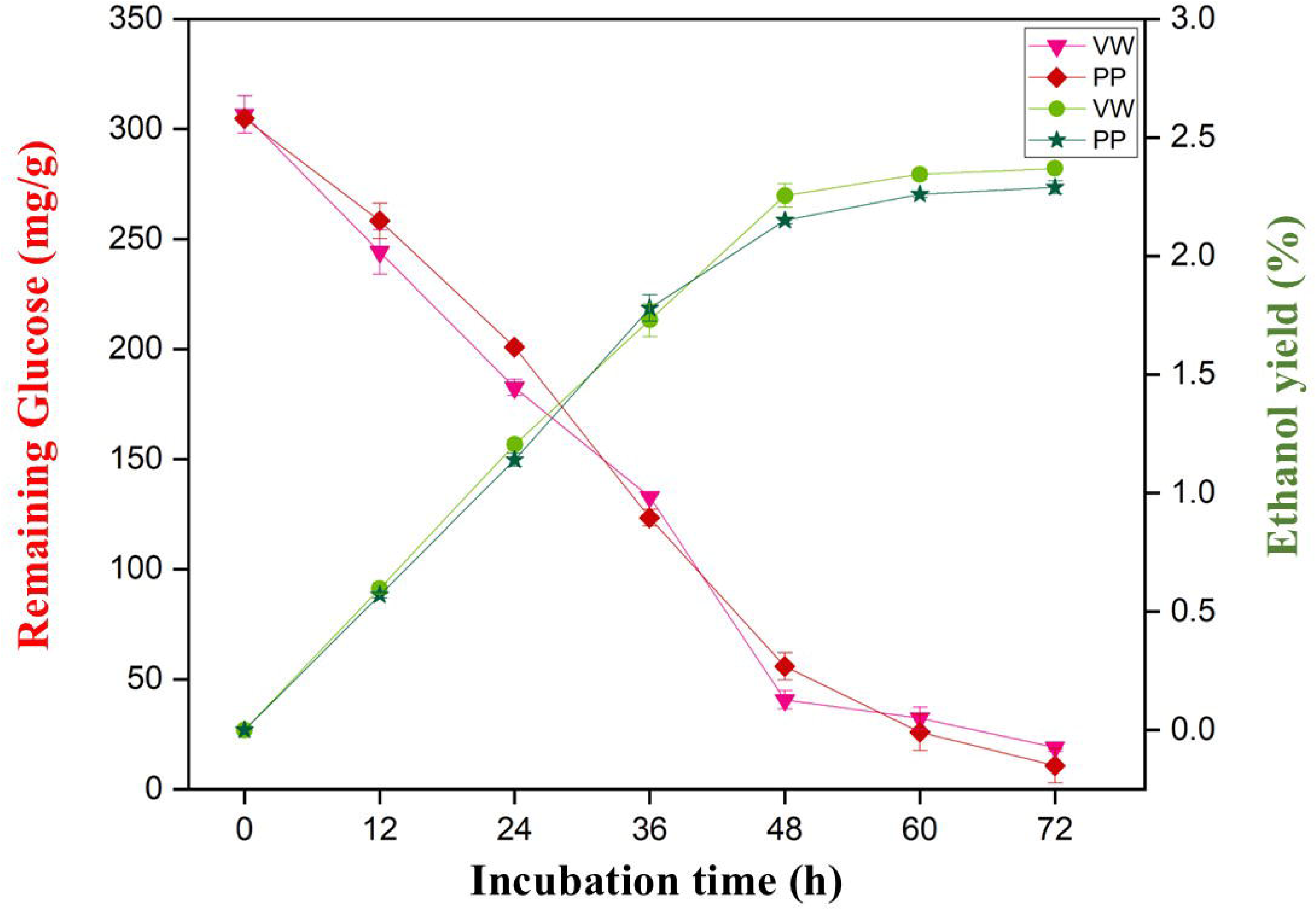
Comparison between consumption of glucose vs. ethanol yield from VW and PP biomasses at multiple time points.

## 4. Discussion

### 4.1. Lignocellulose composition of FVW biomasses: prospects and challenges

An estimated 1.3 billion tons of food is lost or wasted each year during production or processing [42]. In the United States alone, approximately 150 million metric tons (MMT) of food loss and waste have been reported [43]. According to another estimate that vegetable and fruit wastes constitute >50% of food waste in Europe [31]. India is the second largest producer of vegetables in the world after China. A significant proportion of it remain untreated due to the inaccessibility to efficient disposal systems [28]. This results into environmental hazards through emissions of greenhouse gas (e.g. methane due to anaerobic decomposition) and groundwater contamination, while municipalities incur heavy financial liability for waste disposal. As a practical alternative, biomasses obtained from fruit and vegetable wastes (FVWs) could be useful for bioethanol production due to their high carbon sequestration through cellulose and hemicellulose [24, 25, 27]. In contrast to cellulose and hemicellulose, lignin has traditionally been recognized as a deterrent to the cellulosic bioethanol generation, although recent studies highlight its potential to produce useful bioproducts including bioethanol [44, 45].

One major challenge of using high-moisture waste FBWs as a stable bioenergy source is their varying lignocellulose composition that may differ based on seasons and harvesting regions. For instance, the number of hexose carbohydrates (starch and cellulose) detected in composite vegetable, potato waste, sweet potato waste, and yam waste were 33.44%, 57.14%, 53.2% and 58.2%, respectively [24]. However, these assessments may differ if studied LCBs were obtained from another region and season. Indeed, this study demonstrated a large variation in cellulose contents between single and composite FVW and even across bio-replicates of composite FVW due to existing compositional differences. In this study, biochemical cellulose contents ranged from 32.35-47.73%, 15.10-28.23%, 14.88-25.33%, and 15.09-19.20% in VW, FW, OP, and PP, respectively. Another study on four fruit wastes such as acerola fruit bagasse, pupunha peels, pupunha clusters, and mango peels from the Amazon fruit industry demonstrated 25.45-31.30% cellulose, 2.43-23.20% lignin, and 3.53-14.4% hemicellulose content [46].

Such large variation would be less likely in dry, agro-wastes such as sugarcane bagasse, corn stover, rice husk, or rice straw. For instance, one estimate indicated presence of 38.6% cellulose, 15.9% hemicellulose [47], while another study estimated 36.80% cellulose, 19.70% hemicellulose [12]. The biomass of sugarcane bagasse was composed of 56.54% cellulose and 0.97% (w/w) hemicelluloses according to one estimate [25]; and 32-44% cellulose, 27-32% hemicellulose according to another [48]. One major drawback of using these biomasses is the occurrence of silica and higher lignin that interferes with the saccharification reaction. The lignin contents measured in rice husk (16%, [47], and rice straw (19.6- 22.6%, [12, 49, 50] were higher than what was detected by this study in case of FVWs (∼1-16%). Collectively, FVW possesses many good characteristic features for easy and quick conversion to bioenergy to support energy transitions, advance circular economy, and may provide affordable clean energy solution to remote places.

### 4.2. Analytical properties, fermentable sugar and bioethanol yield support the utility of VW and PP for bioethanol production

In this study, analytical methods like ATR-FTIR, TGA, and XRD have been employed to assess effects of various physical and chemical properties of biomasses that may affect the biofuel potential. These methods have previously been applied to other biomasses for effective screening purposes [9, 11]). For instance, the intensity of signature peaks representing cellulose, hemicellulose, and lignin may indicate the total LCB content of the biomass. Similarly, a change in the height of diagnostic FTIR peaks representing cellulose and lignin due to pretreatment is indicative of the efficiency of the reaction (Fig. 3A-D). Several studies demonstrated the removal of different lignin-specific peaks after pretreatment. One such peak at ∼1374 cm^-1^ was removed from rice straw biomass after steam explosion pretreatment [51]. Similarly, two lignin peaks at ∼1594 and ∼1427 cm^-1^ were removed from rice straw tissues after dilute acid pretreatment [52]. A decrease in the intensity of lignin peak (∼1638 cm^-1^) in vegetable waste was also observed after steam and sodium chloride pretreatment [39]. In this study, lignin peaks at ∼1251 cm^-1^, ∼1642 cm^-1^, and ∼1724 cm^-1^ were decreased considerably after pretreatment using 2% NaOH solution. The intensity of peaks indicating cellulose and hemicellulose were also increased after pretreatment. In VW tissues, the intensity of cellulose and hemicellulose peaks at ∼1318 cm^-1^ was increased significantly after steam and sodium chloride pretreatment [39]. The intensity of another polysaccharide peak at ∼2908 cm^-1^ was increased after pretreatment [53]. In this study intensity of cellulose- hemicellulose peaks at ∼3281 cm^-1^, ∼2919 cm^-1^, and ∼1013 cm^-1^ increased significantly, and also new peak representing cellulose and hemicellulose was detected at ∼3347 cm^-1^ (Fig. 3A, B).

Lignin is not the only contributing factor that makes cellulose less digestible to cellulolytic enzymes. Another key determinant is the amount of crystalline cellulose present in the biomass, which deters enzyme efficiency. In a previous study on six grass species, *Saccharum spontaneum* and *Arundo donax* contained the highest CrI (43.63% and 46.96%), which was in contrast to *Sorghum bicolor* (32.51%) and *Coix lacryma-jobi* (33.02%, [11], Table 5). Consequently, the highest glucose yield was recorded for *Sorghum bicolor* (420 mg/g), which was followed by *Coix lacryma-jobi* (263 mg/g). Similarly, in this study, low CrI was observed for VW (19.12%) and PP (19.76%), although another study on composite vegetable waste (CVW) comprised of cabbage, spinach and cauliflower residues obtained higher CrI than this study (63.30%, [39]. This indicates that the composition of the CVW has a big influence on many of its biofuel-related physical properties. In the present analysis, the saccharification reaction demonstrated high glucose yield for VW (325.65 mg/g) and PP (317.17 mg/g). This was comparable to the previous study on CVW, which yielded 302mg/g glucose due to combined use of physical and chemical pretreatments [24] (Table 5). Yield of ethanol from LCB derived sugar can be influenced by fermentation conditions such as efficiency of the microbial strain used, incubation time, pH, temperature, and the initial concentration of sugar [54, 55, 56, 57]. Therefore, the impact of incubation time, pH, sugar concentration was assessed in this study to optimize procedures to transform vegetable waste into bioethanol. The amount of obtained ethanol varied (1.55 - 2.03% for VW, and 1.33 - 1.74% for PP) with a change in pH (6 - 6.9) of the media, but the optimum yield for both VW and PP was observed at pH 6.3 and 72h post incubation. Such impact of pH on ethanol yield has been observed in de-oiled algal biomass, where 0.122 g/g, 0.145 g/g, 0.102 g/g of ethanol was obtained at pH was 5, 5.5 and 6 respectively [58]. Similarly, an optimum temperature (32.2 °C), pH (4.07), and incubation time (72 h) was identified to attain maximum ethanol yield from waste potatoes [57]

**Table 5.** Biochemical comparison of yields with other studies.

| Sample name | Moisture content (%) | Char content (%) | α-cellulose content (%) | Hemicellulose (%) | Acid insoluble lignin (%) | Crystallinity index of cellulose (%) | T<sub>max</sub> (°C) | Glucose yield (mg/g) | Ethanol yield | References |
| --- | --- | --- | --- | --- | --- | --- | --- | --- | --- | --- |
| Assorted vegetable waste | 6.78 | 30.2 | 47.73 | 25.32 | 11.98 | 19.12 | 299.36 | 325.65 | 2.14% | This study |
|  | - | - | - | - | - | - | - | 515.00 | 125.6 mg/g | [24] Chatterjee and Mohan (2021<sup>a</sup>) |
|  | - | - | - | - | - | - | - | - | 14.17% | [55] Promon et al. (2018) |
| Orange peel | 11.77 | 20.8 | 25.33 | 29.15 | 1.706 | 22.18 | 336.66 | 302.98 |  | This study |
|  | 73.53 | - | 69.09 | 5.43 | 19.8 | - | - | - | - | [59] Ayala et al. (2021) |
| Potato peel | 6.5 | 27.2 | 19.20 | 32.43 | 1.116 | 19.76 | 281.66 | 317.17 | 1.69% | This study |
|  | - | - | - | - | - | - | - | 560.00 | 235.4 mg/g | [24] Chatterjee and Mohan (2021<sup>a</sup>) |
|  | - | - | 5.69 | - | - | - | - | - | 6.18% | Chintagunta et al. 2015 [60] |
| Assorted Fruit waste | 8.5 | 24.43 | 28.23 | 27.78 | 6.72 | 24.59 | 317.23 | 247.57 | - | This study |
|  | - | - | - | - | - | - | - | - | 1.18% | Utama et al. 2019 [61] |
| *Sorghum bicolor* | 5.80 | 14.11 | 36.33 | 14.11 | 15.71 | 32.51 | 381.73 | 420.06 |  | Rahaman (2023) |
| *Coix lacryma-jobi* | 6.08 | 19.87 | 41.81 | 15.66 | 16.81 | 33.02 | 366.46 | 263.58 |  | [11] Rahaman (2023) |
| *Bambusa bambos* | 8.60 | 4.12 | 64.53 | - | 21.02 | - | 320.67 | - |  | [9] Biswas et al. (2022) |

## 5. Conclusions

This study aims to explore the valorization of lignocellulosic biomasses of unsegregated vegetable waste as a sustainable and economically viable source for biofuel production. However, future studies may aim to estimate the efficiency of the proposed process in terms of total sugar recovery and ethanol yield. The use of additional enzymes that can hydrolyze pentose sugar such as hemicellulose may allow maximum utilization of the available bioresources. A more cost-effective approach would involve use of combinations (consortia) of microbial strains for delignification, saccharification and fermentation. Collectively, findings of this study may empower rural civic bodies (municipalities) to gain multiple benefits at a time: (i) reduction of green-house gas (methane) emission from FVW biomasses, (ii) expenditure cut on waste management (iii) green energy solution to remote places (iv) employment generation for people involved in the chain from waste collection to bioethanol conversion.

## Credit authorship contribution statement

Sudeshna Bera: Methodology, Data analysis, Writing - Original Draft Sakhi Kundu: Methodology, Writing - Original Draft

Sarthak Mondal: Methodology, Writing - Original Draft Semantee Bhattacharyya: Methodology, Writing - Original Draft Asmita Banerjee: Methodology, Writing - Original Draft

Asmita Das: Methodology, Writing - Original Draft Touhidur Rahaman: Data analysis, Writing - Original Draft Pijus Ghorai: Data interpretation

Sukanta De: Methodology and data interpretation

Jhuma Ganguly: Data interpretation, Writing - Original Draft

Avishek Banik: Methodology (Fermentation) and data interpretation

Malay Das: Funding acquisition, Supervision, Project conceptualization, Writing - Review and Editing

## Declaration of generative AI and AI-assisted technologies in the writing process

The authors declare that During the preparation of this work the author(s) did not use any AI and AI-assisted technologies.

## Data availability

Data will be made available on request.

## Acknowledgement

Research results reported in this manuscript have been funded by theYouth for Undertaking Value Added Innovative Translational Research (E-YUVA) scheme of BIRAC, DBT (BT/EF0099/01/22) and FRPDF grant of Presidency University. Research fellowship was provided to TR by the UGC MANF and to SB by UGC-JRF. The authors thank Dr. Biplab Maji and Ayanjyoti for helping with TGA data. We acknowledge UGC-DAE CSR, Kolkata Centre for XRD sample run.

---

Fig. S1 Identification of diagnostic glucose specific peaks identified in the four FVW tissues; OP (A), PP (B), FW (C), VW (D) by observing the diagnostic mass fragmentation pattern in the High-Resolution Mass Spectrometry (HRMS) analysis. Abbreviations used: FW- fruit waste, OP- orange peel, PP- potato peel, VW- vegetable waste.

