## Supplementary figures and images for "Comparative evaluation on the fermentative potential of single and assorted fruit and vegetable waste for production of bioethanol"

### Supplemental Figure 1

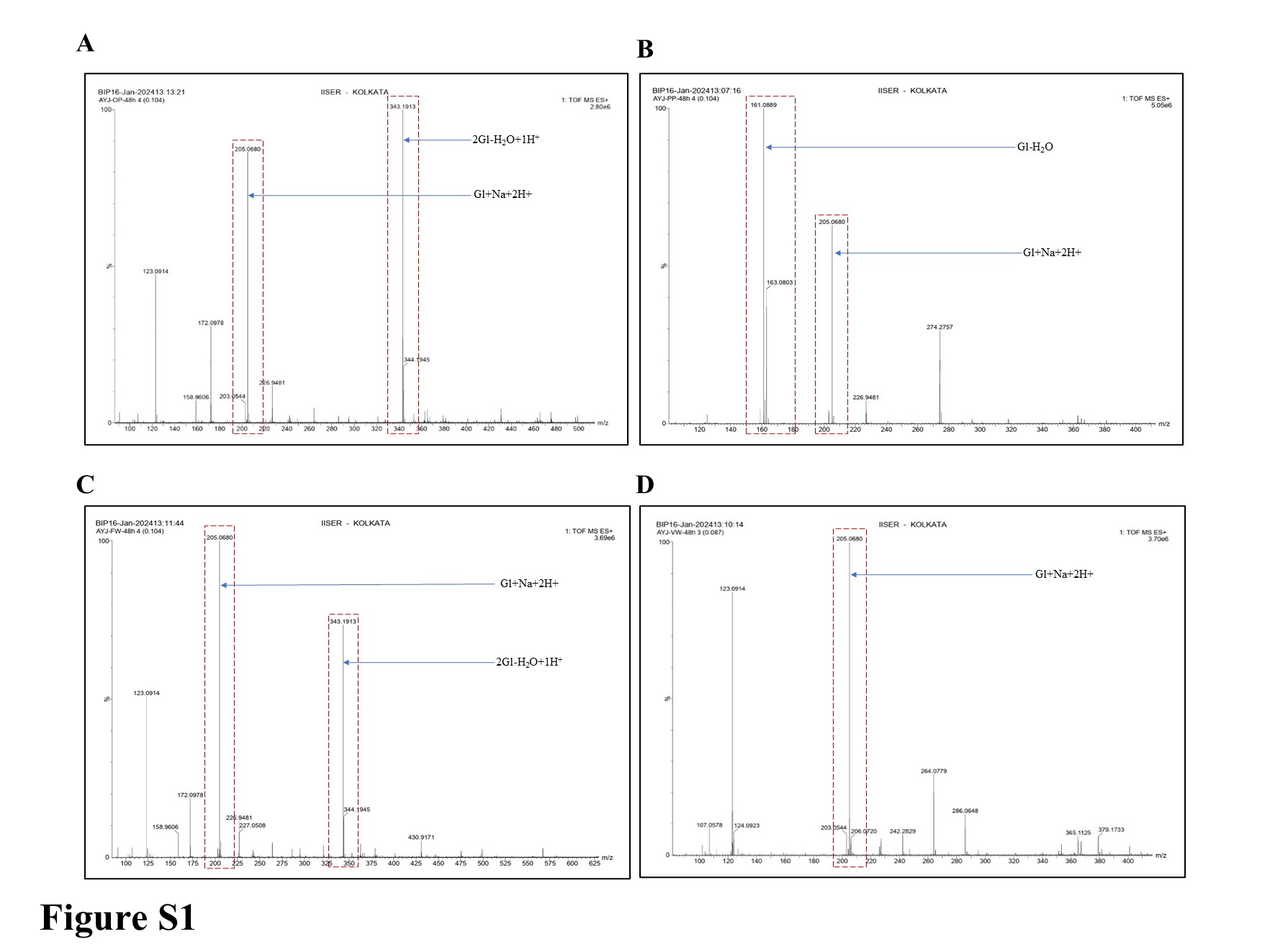
